# Listening to children’s and educators’ experiences in long-term place-based conservation education sessions on southern Indian rocky outcrops

**DOI:** 10.1101/2025.08.05.664756

**Authors:** Vijayan Jithin, Rohit Naniwadekar

## Abstract

Interactions with nature can influence people’s perceptions of nature and its conservation. Connected to this, there is an increasing research interest in improving childhood nature experiences and environmental awareness through direct outdoor educational activities. While this research largely remains geographically biassed to the nature of temperate regions, global and national policies are pushing towards place-based outdoor environmental education in other geographies where research and theory-practice gap persists. The existing environmental outdoor education literature identifies the on-ground challenges, mostly from educators’ perspectives and focusing on academic achievements of participants. A better understanding of culturally specific, locally appropriate, and diverse pedagogies, drawing on both children’s and educators’ voices and relational values is required to fill this gap. We studied ∼20 years old, unique place-based conservation education sessions on the rock outcrops in Kannur (Kerala, India), facilitated by schools and non-governmental organizations. These sessions introduce children to the experience, ecology and conservation issues of rocky outcrops, an easily accessible local socio-ecological system. Considering this as a case-study, using mixed-methods, we aimed to assess (i) if and how these outdoor sessions improve socio-scientific understanding, and positive attitudes towards these habitats and their conservation in children, and to (ii) identify the sessions’ strengths, challenges, and relational values reflected in them through long-term experiences of its participants and educators. Our within-participant pre- and post-outdoor session surveys showed quantitative evidence for short-term increase in overall knowledge, but not any notable changes in measured attitudes or interests. But, the qualitative analysis of in-depth interviews with children and educators revealed major strengths of the sessions around the Natureculture elements, and relational values; while the challenges were mostly around educators’ understanding of children, pedagogical design, and logistics. Drawing on the long-term experiences of educators and children in the backdrop of existing global literature on outdoor education, we provide new insights on how using the power of emotions and experiences, positive and solution-oriented outlooks, unique socio-ecological elements, and relational values can be beneficial in designing conservation education sessions in often overlooked geographic and cultural contexts.

## INTRODUCTION

There is increasing evidence for declining human–nature interactions, potentially impacting human well-being and people’s support for biodiversity conservation (Soga & Gaston, 2016; 2020). These interactions, which could be positive and negative, can influence people’s attitudes and behaviour towards nature. A better understanding of these contrasting experiences are crucial in addressing the complex conservation challenges (Evans et al., 2023). Towards this, there is an increasing research interest in improving childhood nature experiences through direct outdoor educational activities such as nature camps and field trips (Ardoin et al., 2020; Mann et al., 2022a). This geographically-biased research interest, mainly from the nature of temperate regions, is based on studies suggesting early childhood as a crucial time for developing environmental literacy and pro-biodiversity behaviours in adulthood, in connection with the sociocultural learning theory (Ardoin & Bowers, 2020; Evans et al., 2023; Aota & Soga, 2024). Many studies from these geographies show that such educational activities positively impact cognitive, affective, interpersonal, social, physical health and behavioural domains of children (Ardoin et al., 2020). Previous Environmental Education (EE) and Outdoor Learning studies also show that childhood nature experience can provide important learning opportunities, and improve environmental attitudes and academic achievements (Rickinson et al., 2004; Wu et al., 2023; Beery & Jørgensen, 2016). Such experiences can also improve socio-environmental consciousness when contextualized within place-based, local conservation challenges (Hohenthal & Veintie, 2022; Varela-Losada et al., 2016; Häyrynen et al., 2021).

Given the policy-level focus on EE to improve biodiversity conservation at global- and national-levels (Ardoin et al., 2020), and our geographically-biassed research landscape (Evans et al. 2023; Yemini et al., 2023), it is important to understand how participants and educators are experiencing these place-based EE sessions in largely neglected contexts. Also, our understanding of the conditions under which particular outdoor learning approaches are most effective for various desired outcomes (socio-emotional, academic and wellbeing benefits) remains limited (Mann et al., 2022a). For example, how do we move beyond the typical dichotomic focus on instrumental (value for a person) and intrinsic (independent of human valuation) values of nature in EE, and harness the power of relational values, pertaining to all kinds of relationships between people and nature (Chan et al., 2016; dos Santos & Gould, 2018)? This approach could be particularly useful in addressing real-world pedagogical challenges within unexplored socio-ecological settings, especially where nature, culture, society, and humanity are not viewed as separate entities (Eyster et al., 2023; Reid et al., 2021; Himes et al., 2024). Furthermore, assessment frameworks for EE sessions are largely based on adults’ perspectives, and not on the experience of participants, resulting in exclusion of nuanced considerations from children’s perspectives (Giusti et al., 2018; MacQuarrie, 2022; Marchant et al., 2019). Also, one of the key challenges identified by multiple studies is the time constraint imposed by the existing academic curriculum (Yemini et al., 2023; Marchant et al., 2019). This often necessitates schools and non-governmental organizations (NGOs) to rely on short-duration outdoor experiences, over the longer ones.

For example, in India, where EE is a regular school subject, recent educational policies are pushing towards formal place-based outdoor EE (NSCNCF, 2023; SCERT, 2023). As EE’s key concepts of environment and literacy are culturally specific, to implement such policies, there is a need to understand pedagogies that are locally appropriate, rooted in easily accessible ecosystems (than protected areas), diverse, and inclusive – which can address the research and theory-practice gap in these geographies (Jithin et al., *in press;* Almeida & Cutter-Mackenzie, 2011; Cole, 2007). To achieve this, it would be beneficial to investigate the long-term experiences of stakeholders already involved in such sessions, particularly in poorly understood ecosystems, beyond the commonly studied forests and parks. India has a few such grassroots initiatives offering their long-term experiences.

This study synthesises the long-term experiences from place-based conservation education sessions on the rock outcrops in Kerala, India, which include formal and informal sessions facilitated by schools and NGOs. These short-term (1-2 days) sessions for children provide direct nature experience, and introduce them to the socio-ecology and conservation issues of lateritic rock outcrops. The pioneering educational sessions were catalysed by the protests led by local people against the china clay and lignite mining, but later on continued by the local organizations (Anthony, 1996; Unnikrishnan, 2024). These outcrops face multiple threats including land-use conversion, mining, and tourism, while harbouring endemic and threatened biodiversity, and unique cultural values (Jithin et al., 2024; KFRI, 2019; Watve & Chavan, 2020). These ecosystems are officially classified as ‘wastelands’ in India, though being one of the most threatened among tropical habitats, and despite their potential to be used as educational tools beyond Geo-education (Wolniewicz, 2021; Michael & Lindenmayer, 2018; Watve, 2013).

Considering this as a case-study, we carried out a mixed-method research covering the organizers, educators, and participants of these outcrop-based outdoor sessions, using quantitative surveys and in-depth interviews, followed by generating metainferences (Stern et al., 2014; Younas et al., 2023). In a pre-post test framework, we quantitatively assessed if and how one such half-day session can influence participants’ socio-scientific understanding of the rock outcrops and attitude towards its conservation; and documented participants’ unique session experiences. In the second phase, using interviews, we qualitatively explored children’s and educators’ learning, experiences and perspectives on the strengths, challenges and key relational values reflected in sessions occurring across Kannur. Given the influence of outdoor education sessions in improving students’ knowledge, attitudes and beliefs (Bradley et al., 1999; Moseley et al., 2019), we expected an increase in children’s formal knowledge, and pro-environmental attitudes related to rocky outcrops after these sessions. Though many previous studies mainly focus on adults’ perspectives, equal importance was given to children’s voices in our study, since learning frameworks can greatly benefit from it by bringing in nuanced features. Throughout the study, we viewed children as competent social agents and co-constructors of their own worlds, whose views provide rich, reliable descriptions, definitions, and examples of their reasons (Spiteri, 2021; Barblett et al., 2024).

## MATERIALS AND METHODS

### a. Study area and context

The study was conducted in two phases, covering multiple lateritic plateaus in Kannur, Kerala, India. In the first phase, we piloted our questionnaire in partnership with the Madayi regional committee of Kerala Sasthra Sahithya Parishad (KSSP), a people’s science movement in Kerala (Heller, 2001), during their session. The original survey followed this, in collaboration with a government school in Kannur, and the second phase consisted of interviews with individuals associated with three different NGOs and seven schools in various locations. The first phase was exclusively carried out in Madayipara, a prominent flat-topped lateritic hillock in Kerala harbouring unique, endemic and threatened biodiversity, and home to multiple archaeological remains, a sacred grove, and many cultural and religious structures (Bhagyalakshmi, 2024; Palot & Radhakrishnan, 2005; Fig. 1). Different NGOs and schools conduct yearly nature camps on Madayipara, some having history dating back to more than 20 years (Balakrishnan & Jafer Palot, pers. comm.; Anonymous, 2011).

**Figure 1.**
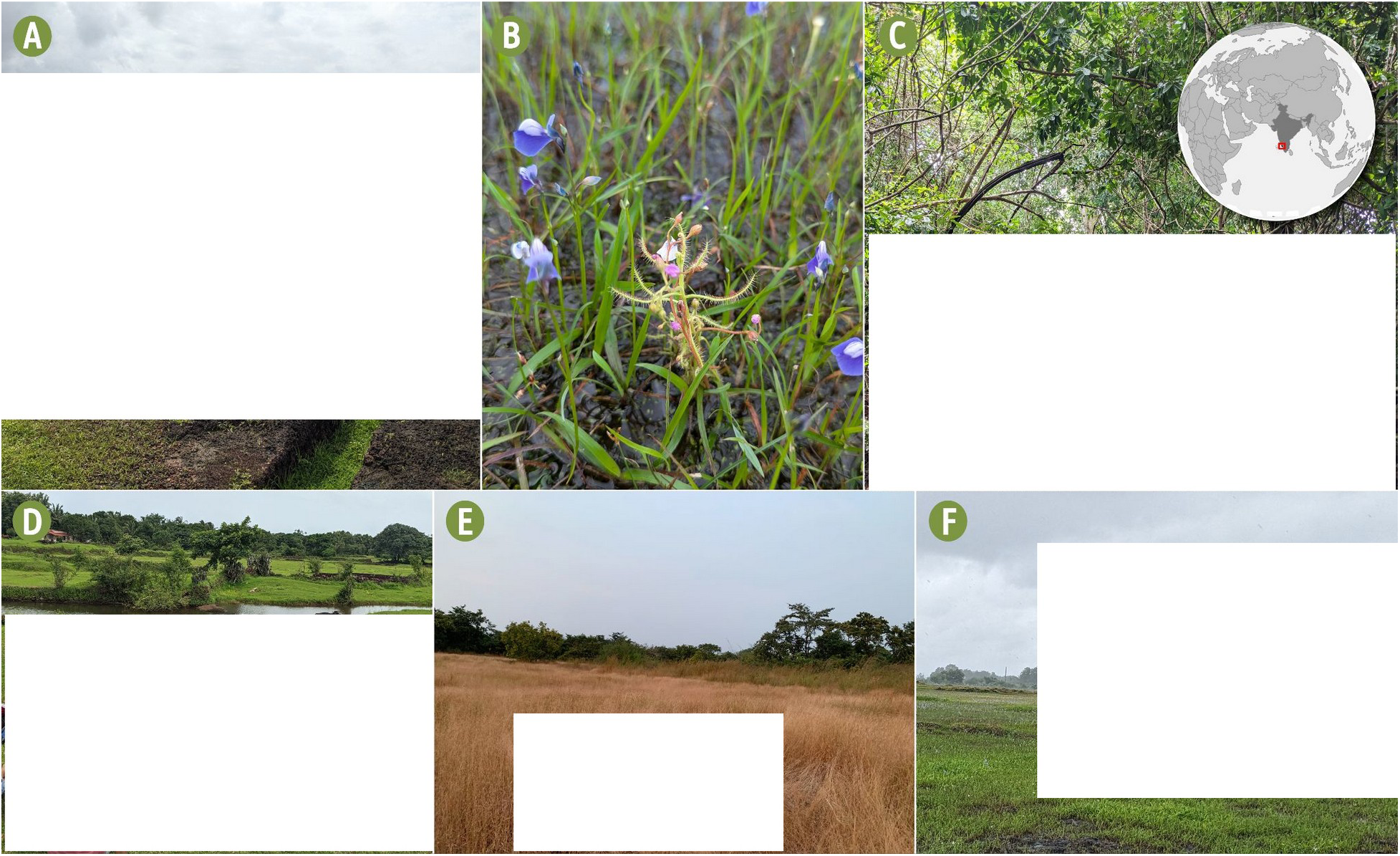
Photographs showing (A) participants of the conservation education sessions exploring the open plateau; (B) ephemeral plants including *Drosera* sp. and *Utricularia* sp. on the rock surface; (C) inside the sacred grove, and (D) near a historical pond; experiencing (E) rocky plateau evenings in summer and (F) rainfall during monsoon. Photographs by Saaji Iringal, Anaswar K, and Jithin Vijayan. Individuals featured in the photographs have given their permission. Inset orthographic map shows the study region - Kannur, Kerala, India. *Note: portions showing identifying information of people have been removed from photographs, in compliance with the policy of bioRxiv*.

Madayipara is surrounded by various wetlands and rivers, and is a unique ‘amphibious’ ecosystem with different wet and dry phases. This 365 ha area is home to 636 plant species with many narrow endemic and rare species that are only present in the area (Pramod & Pradeep, 2021), and 255 species of birds, with 110 migrants, illustrating the biodiversity value of the area (Palot et al., 2020). The camps and field trips to Madayipara largely follow walking pedagogy (Friedman et al., 2025), and generally involve exploring different habitats including the sacred grove called ‘Madayi Kavu’, large rock pools, archaeologically important Jewish Pond and fort remains, and often involving getting wet in the rain on the rocks during the monsoon period.

### b. Study framework

In the first phase, we pilot tested our self-reported survey questionnaire (N = 34; Appendix S1) during the ‘rain experience camp’ by KSSP in July 2024, using a within-subject pre- and post-test design. The questionnaire contained multiple choice, likert-scales, and open-ended items relevant to socio-scientific understanding of the rock outcrops, spanning across multiple values of nature, participants’ perceptions and experiences (Jithin & Naniwadekar, 2025; Gould et al., 2018). Following this, adolescents between the ages of 15-18 participating in a field trip to Madayipara in October 2024 were recruited from a government school within a 25 km radius of Madayipara. They were National Service Scheme volunteers from different subject backgrounds, studying in the 11^th^ and 12^th^ grades. Similar to the pilot-test, a pre-session instrument was administered guided by the researcher at the beginning of the field trip, by reading aloud each item and giving students sufficient time to answer (Larson et al., 2011; Appendix S1). This session was guided by one experienced educator, and the researcher (VJ), at the request of this educator, leading to active participation of the researcher in the session, which mainly was educator-led and included intentional teaching (Speldewinde, 2022; 2024). Following the session, which lasted between 11:00 AM and 02:00 PM, the post-session questionnaire was administered after a week, at their school. Among the 47 individuals, only 43 filled both pre- and post-session questionnaires, limiting further analysis to these participants (28 girls, 13 boys, 2 gender not disclosed), except otherwise noted.

During the second phase (19 September 2024 to 28 February 2025), semi-structured interviews were conducted with participants, educators and organizers of the short-term nature education sessions based out of rocky outcrops, using a snowball sampling approach. This included the *Mazha Nanayal Camp* (Rain experience camps) by KSSP at Madayipara and its satellite version in Sreekandapuram, *Pookkaala Sahavaasa Camp* (Flowering season cohabitation camp) jointly organized by the Society for Environmental Education in Kerala (SEEK) and Malabar Natural History Society (MNHS), and sessions from government and government-aided schools in the region. Many interviewees’ experiences and memories overlapped between these different types of sessions, leading us to focus on the common aspects of these sessions, rather than the differences. Many organizers among the interviewees themselves were educators, and vice versa. Henceforth, the term ‘educators’ represents both educators and organisers in the following text. It is also important to note that the educators interviewed may have been influenced by their own experiences as participants in similar sessions during their childhood. The interview sample consisted of 23 participants aged 11-16 years (13 girls and 10 boys; 2021-2024 sessions), three previous camp participants aged 24-32 years (one women and two men; 2003-2016 sessions) and 15 educators aged 40-67 years (three women and 12 men; 1999-2024 sessions; Appendix S2). Due to logistical constraints, in three instances, the interviewer explicitly interacted with children in groups, and there were instances where children came in between interviews and responded to questions for other participants. We considered all these responses for the analyses. Before the interviews, the interviewer (VJ) ensured rapport was established. All children were interviewed in the presence of parents or their teachers, either at their home or school, adapting to their conversation styles. Interviews were conducted in *Malayalam*, the native language in the region, and audiotaped with permission, for clear transcription and translation. Interviews generally lasted between 30-100 minutes, with breaks and other activities adapting to situations.

All questionnaires were prepared based on observations from previous literature, and educators (Salazar et al., 2021; Krosnick & Presser, 2010), and reviewed by the Ethics committee, school teachers, and experienced researchers beforehand. VJ, a native speaker of *Malayalam*, handled the translation of responses to English, ensuring matching both language versions across translations for both the questionnaire and interviews (Brislin, 1970). Ethnographic notes were taken during all occasions.

The data collection procedures and practices were approved by the Nature Conservation Foundation Research Ethics Committee (NCF-EC-23/07/2024-(92)), and by the school authority (NSS/45/2024-25 dated 9 October 2024). Participants were informed about the research purpose, voluntary nature of the study, benefits and data confidentiality prior to data collection, and written consents were obtained. In addition to this, before the pilot KSSP session, we explained about the study to the organizers and parents in an online session. Prior to each interview with children, the researcher contacted parents or teachers to obtain consent to visit the sites, and interact with children.

### c. Quantitative methodology

The survey questionnaire administered before and after the field trip evaluated the change in participants’ (i) socio-scientific understanding of the lateritic plateaus, (ii) positive and negative attitudes towards the plateau conservation, (iii) session experience, and (iv) attitudes to environmental conservation. In addition to this, socio-demographic data concerning age, gender, locality, and previous visit to the area was also obtained. The score for each question concerning socio-scientific understanding of the lateritic plateaus was summed to calculate a composite ‘plateau knowledge score’ for each participant. This score was compared before and after the session, using a Wilcoxon signed rank test with continuity correction since the data was not normally distributed, and ‘coin’ package was used to calculate the effect size. In addition to this, we examined score changes for different aspects (e.g., cultural, historical, animals, plants) after the session.

For the attitude-related likert-scale questions, Cumulative Link Mixed Models (CLMMs) were fitted with ‘probit’ links to examine the effect of the session (fixed effect), after accounting for individual level differences (random effect), using the ‘ordinal’ package (Taylor et al., 2023; Christensen et al., 2023). Fisher’s exact test was used to assess the participants’ change in interests. Open-ended questions in the survey instruments were inductively coded to categorise the response and frequencies were analysed. All analyses were carried out in R (v4.4.3; R Core Team, 2025).

### d. Qualitative methodology

In the case of participants, (i) learning and (ii) experiences from, and (iii) suggestions to improve these sessions were explored. From educators, we explored their (i) aims, objectives and motivations, (ii) preparations, (iii) challenges concerning the sessions, and (iv) changes over the years, and (v) suggestions to improve the sessions.

For the content analysis (Braun & Clarke, 2006), transcribed interview scripts in English were first tagged with pre-decided question themes in Taguette (v1.4.1) web-application hosted offline (Rampin & Rampin, 2021). The exported spreadsheet containing tags from all interview scripts was then subjected to detailed inductive coding in LibreOffice Calc (v7.5). VJ coded the interview scripts within and between categories by repeated classification, merging, and reconstruction to identify primary patterns of themes and subthemes (Fig. S4). The original codes were further revised considering the outliers, contrasting perspectives, and emerging new themes. The main coder (VJ) is from a lateritic plateau landscape in another location in Kerala, and have participated in programmes of KSSP and MNHS in the past, but not the ones studied in this work, which should be acknowledged in thematic analysis (Braun & Clarke, 2019). Individual researcher positionalities were acknowledged and assumptions were questioned in regular meetings, and with our collaborators. To enhance the clarity of resulting ideas, we have reported quotations, with a few grammatical corrections, to retain the interviewees’ original thoughts.

## RESULTS

### a. Phase I - Insights from the written responses

Among the field trip participants, 81.4% had already visited Madayipara before the session. Participants reported that seeing the water sources (e.g., sea, river, stream, rock pools) and nearby viewpoints, and discussions on flowers and sacred grove were the most interesting parts of the session. While 38 (88.4%) participants liked the session, one disliked it and three gave a neutral response.

The overall plateau knowledge score was significantly higher after the session (*V* = 131, *p* < 0.001, pseudo-median = -3.5, 95% CI [-4.5, -1.5], *Z* = 3.60, *r* = 0.548; Fig. S1). Scores for plateaus’ cultural aspects, identification of flora, formation, ecosystem knowledge, and uniqueness showed a marked increase (Fig. 2). However, there was only a slight increase in the knowledge about fauna, historical aspects, and place knowledge. Identification of threats had a comparatively greater proportion of extreme increase and decrease in scores (Fig. 2). There was no score change for 33-60% of participants across different aspects.

**Figure 2.**
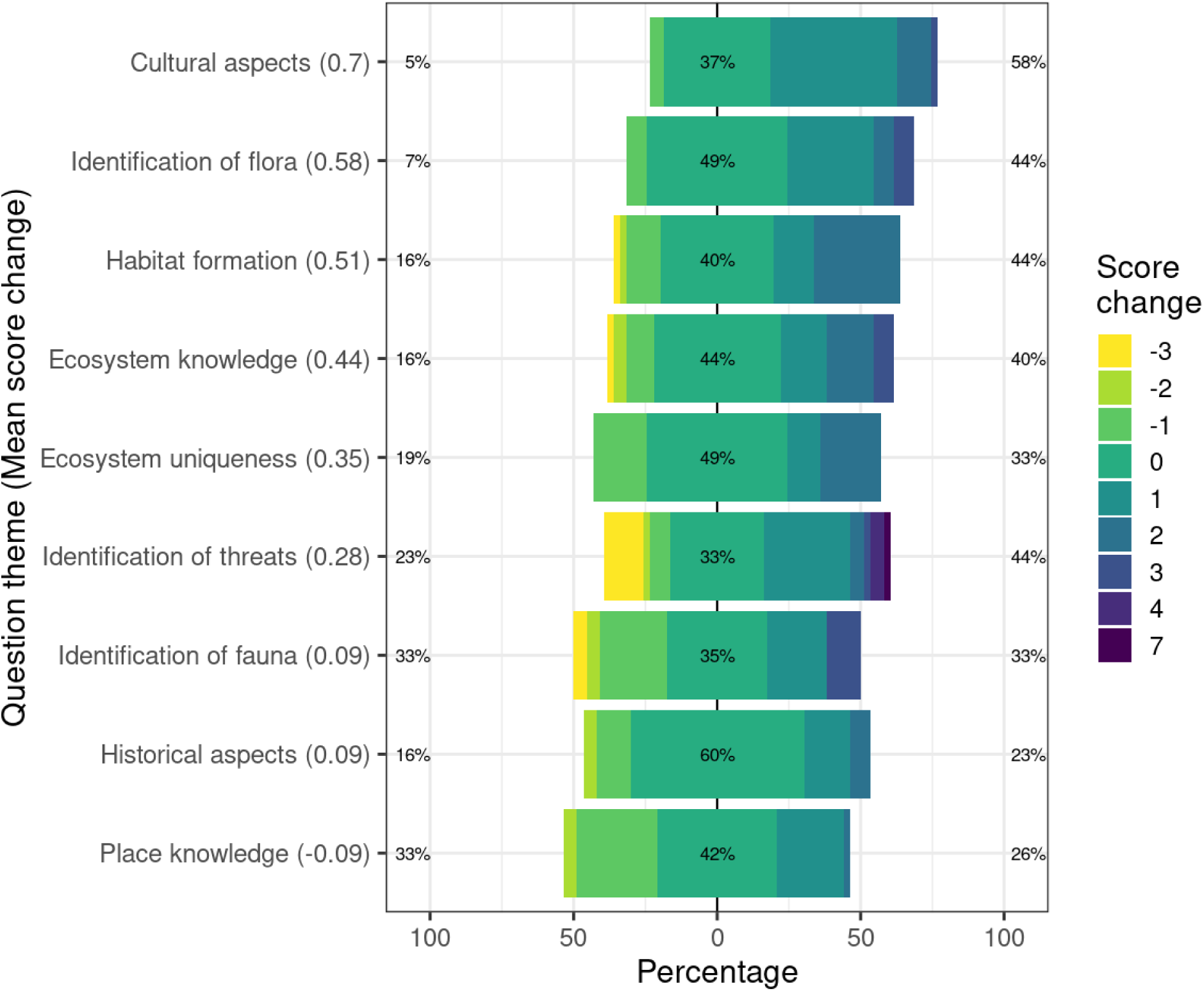
Diverging chart showing the score change after the session for the knowledge-based questions on different themes. The area of different colours (score change categories) is proportional to the number of participants in that category. The percent numbers labelled on the zero category at the center indicate the proportion of participants without any change in their scores after the session. The percentage labels on the left and right axes show the proportion of participants with a decrease, and an increase in their scores, respectively. The numbers in the parentheses to each question theme indicate the mean overall score change for all participants.

Prior to the session, most participants reasoned that the prevalence of threats would result in loss of nature, beauty, tourism opportunities, and serenity in the plateau. After the session, more participants suggested that threats would result in increased pollution and loss of nature. They also suggested a need for living in harmony with nature and a need for increased research and educational activities (Fig. S2A-a).

Participants’ reactions to the news featuring the protest against quarrying on the plateau revealed contrasting responses and ambivalence (e.g., job opportunities vs. environmental costs) (Fig. S2A-b). During both pre- and post-sessions, 59.1% of the participants identified problems posed by quarries for the plateau. There was a marginal increase in participants who explicitly supported the protest from 27.3% (pre-session) to 29.6% (post-session) (N = 44).

> *“It is harmful for the environment. Quarry workers are living off of this job. So, it is difficult for them.”*-(R9; Female, 17 years)

> *“Why do we need quarries? Why destroy a beautiful, natural place? People’s reaction is very correct. Can protests alone bring change? Don’t governments need to acquire similar places?”*-(R43; Female, 15 years)

Participants who were ambivalent about quarrying reduced from 15.9% to 9.1%, and only a few requested more information on it after the session (15.9% to 6.8%). One individual expressed concern for people dependent on quarries for livelihood during the pre- and post-session. There was an increase in the respondents questioning the benefit of destroying nature (2.3% to 18.2%), and questioning the efficacy of protests against quarrying for preserving the plateau (2.3% to 6.8%). Before the session, participants mentioned the negative impacts of quarrying on nature, its aesthetic beauty, on their ‘native’ place (*Naadu*), and peoples’ lives (e.g., water availability, etc.). After the session, participants mostly mentioned the need for preserving nature and their native place from quarrying.

Overall, more than 80% of participants were interested in seeing *Drosera* sp., (positive experience) an insectivorous plant found on the plateau, consistently before and after the session. Compared to this, only 35% were interested in seeing scorpions (negative experience). After the session, we failed to detect statistically significant change in the participants’ interest to see *Drosera* or scorpions (*Drosera*: *p* = 0.5477, *OR* = 0.58, 95% CI = [0.13, 2.23], Scorpion: *p* = 0.5016, *OR* = 1.41, 95% CI = [0.53, 3.8]). But, their motivations to see the organisms differed after the session. Participants reasoned novelty as the main motivation to see *Drosera* prior to the session, while they mentioned its uniqueness, beauty, rarity, and their curiosity to see its prey-trapping process during post-session. During the pre-session, the participants mentioned that there was no novelty in observing a scorpion (as most participants had seen one before), but post the session they mentioned scorpion as a ‘dangerous’ animal and they do not like it (Fig. S2B).

There was no statistically significant difference in participants’ opinion on the need for special protection of plateau biodiversity (Fig. 3A; Table S1), or their ability to solve environmental problems (Fig. 3B; Table S1) or the proportion of participants who thought ‘wasteland’ classification is appropriate (Fig. 3C; Table S1) or those who liked outdoor explorations (Fig. 3D; Table S1). In both pre and post surveys, more than 95% participants consistently agreed they will ‘enjoy their lessons more if they learn about similar organisms and habitats in schools, while only less than 10% of the total participants opined it will increase the difficulty in learning (N = 42).

**Figure 3.**
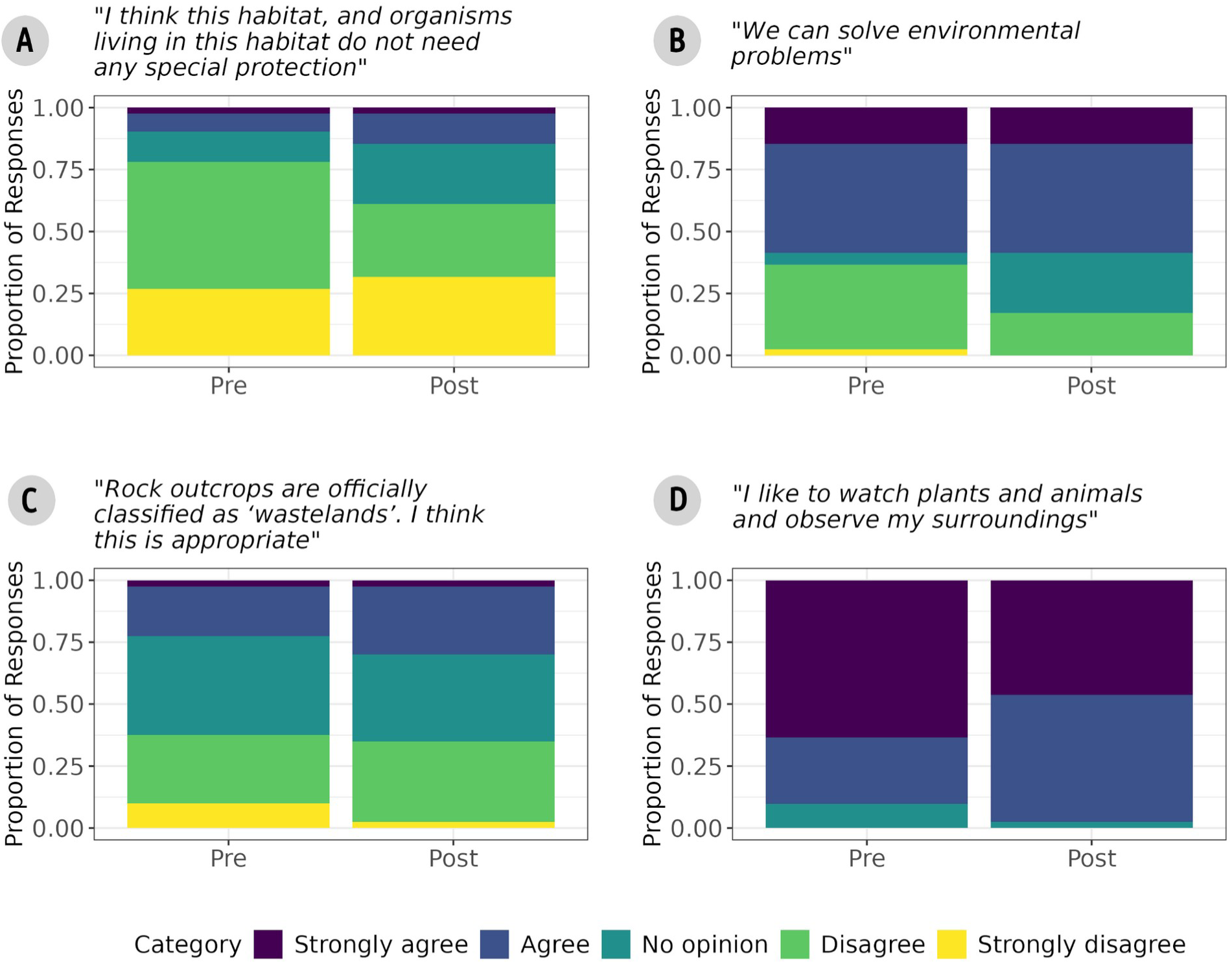
Participants’ response before and after the session to the attitude related statements concerning the conservation of rocky outcrops, hope, and interest in outdoor exploration. None of the changes were statistically significant.

### b. Phase II - Insights from interviews with participants

#### i. **Learning and experience from the sessions**

The sessions provided knowledge on geography, environment, and conservation issues to the participants, and it also helped the learning process in general (Fig. S3-1). This included understanding of the flora, fauna, physical geography, and ecological concepts (e.g., food web, ecosystem services). Knowledge on human geography reflected quite frequently when they talked about the local history, culture (e.g. *Maritheyyam*, an ancient ritualistic performance associated with Madayi sacred grove), local folklore and myths (e.g. crocodiles used to live in the rock pools), and the area’s importance. The environmental history of Madayipara including the china clay mining and protests often reflected in participants’ memories.

> *“When there was a China clay [quarry], households that were downstream to the quarry did not get water. Then the villagers came together, protested, and it [quarrying] stopped. Then all the households [around the quarry] got a good amount of water, even during summer. That is good for people.”* - (P3; Male; 12 y.o.)

Participants were able to list various conservation issues of rocky plateaus, their consequences, and possible solutions. While the participants were mainly focused on tourism and development issues, they were able to think from multiple perspectives. Some students were uncertain about how to strike a balance between conservation and development. Given the negative human impacts on nature, some children were deeply worried about their agency, and a better future for themselves.

> *“Probably [the plateau] will not exist [in future]. They are making roads and portions [of the plateau] are already gone. If something like a government project comes, what can we do? Even if we tell [someone], we are children, right? Don’t know if it will be effective. If there is an opportunity, we will try for sure. Slowly we are destroying nature, though our future is at stake.”* - (P8; Female, 15 y.o.)

Participants found that the knowledge gained during these sessions on rock outcrops complemented the classroom activities and examinations at schools, and helped in extracurricular activities such as educational camps outside schools. These sessions also helped the participants in improving imagination, learning skills, independent thinking, and the ability to explain concepts to peers.

> *“If we destroy them [lateritic plateaus], organisms living there will also get destroyed, right? For example, birds that lay eggs on the rocks will have specific temperature requirements for successful reproduction, right?”* - (P8; Female, 15 y.o.)

Participants’ session experience centered on location-specific (within outcrop) features and experience styles (Fig. S3-2). Beauty of the area, rare and unique features of biodiversity and history were prominent in the location-specific experiences. Hands-on activities, interaction with peers, and exposure to new knowledge appeared as unique experience styles. Many participants particularly enjoyed the outdoor rain experience and the water seepage in lateritic caves.

> *“I remember the cave quite clearly; it was a really nice experience. The water drops were falling from above. It is great that the precious water is being stored in the rocks.”* - (P7; Male, 24 y.o.)

#### ii. Suggestions to improve the sessions

Some participants suggested logistics, experience, content, and teaching-style related improvements, while others were satisfied with the existing session formats (Fig. S3-3). The suggestions included having more guides for outdoor explorations, a more relaxed schedule with lower information load, more games and unstructured time, opportunities for participants to lead sessions, expanding the participant pool to beyond Madayipara and nearby regions for more interactions, and better promotion of the event. Despite acknowledging that weather is unpredictable, lack of rain experience was a main concern for many participants. Participants desired more engaging and curiosity-inducing content that would expand upon their current understanding. They suggested that the sessions be more integrative instead of the current structure of the sessions which is organised in discrete topics (i.e., plants, animals, history, and conservation). While some desired more content on plants, birds and history, others were satisfied with the existing content. Participants also noted a lack of discussions on ‘positive’ or ‘hopeful’ conservation issues.

### c. Phase II - Insights from interviews with educators

#### i. **Aims, motivations and preparations**

While certain educators indicated that the sessions had clear aims, others felt that camp’s purpose was purely experiential (Fig. S3-4). Educators who advocated an experiential format described nature as an open book. Direct and structured aims as pointed out by others were learnings on conservation of lesser-known biodiversity, environment and water, academic concepts helpful in schools (eg. insectivorous plants, concept of ecosystem), and place-based knowledge on laterite rocks, history, and environment. Some educators also pointed out a few indirect aims. This included fostering socialization, spark curiosity, maximize direct outdoor childhood experiences, strengthen nature connectedness through eco-spiritual and ecocentric love. It is noteworthy that all educators opine their aims are fulfilled in the sessions.

> *“Organizers ask [us] to explain to students whatever we see there. They also wonder if we can turn that observation-based knowledge into love; to make children understand what happens to nature if it [an organism, or interaction] is destroyed. This could spark a lifelong interest and love [towards it] in children. When they revisit this place, they will recall these observations. This slowly fosters a gradual interest in their surroundings—a key lesson in conservation.”* - (E2; Male, 49 y.o.)

The motivations of educators to take part in the sessions included their intention to create compassionate individuals for the future, and spread conservation awareness through experiences and emotions (Fig. S3-5). These were inspired by their own past experiences as participants in similar sessions, place attachment, general interest in volunteering, and teaching or interacting with children. Some educators prepared for organizing, and designing the session’s content and style, though others did not specifically prepare (Fig. S3-6).

#### ii. Educators’ challenges, and proposed suggestions

Educators identified many challenges in conducting the sessions, and suggested solutions to some of them (Fig. S3-8;9). Content-related challenges included the dynamic nature of outdoor contexts, difficulties in topic selection and updates, and a lack of authentic historical and cultural resources. Existing educational materials with pre-written answers for curricular outdoor activities, lack of focus on non-academic, direct emotional experiences also contributed to this. Educators acknowledged their ‘conservation vs. development’ dilemma impacting the work, and lack of in-depth (long-term, behavioural) evaluations of the session outcomes.

> *“There was something that perplexed me during these regular plateau trips. In one house, the elder girl was getting married, and the parents wanted to sell a portion of their plateau land, due to financial hardships. But her younger sister, studying in 7^th^ grade in our school did not allow it, saying not even death will deter her. Here, I was the reason…and the parents called me. Given the situation, with whom should I stand?… Eventually at the end, somehow, she let them sell a portion of land. So, children’s minds are falling for this, maybe not all.”* - (E14; Male, 52 y.o.)

Challenges around dealing with children were much more detailed and nuanced (Fig. S3-8). For educators, it was challenging to understand the backgrounds, strengths, and individual interests of all the participants, especially in multi-age-group sessions. Some educators assume children ‘do not know many things’, gain little from sessions beyond recreation, and require moral development lessons. However, others disagreed with this view. While some educators believed children can socialize independently, others emphasized the importance of team-building activities and purposeful games for achieving this. Also, to some educators, all children may not be skilled in persistent and unguided explorations and collective inquiry, while others disagreed, and suggested more guides to facilitate guided or independent exploration by children. To better understand children, some educators highlighted pre-session considerations, including streamlining registration and gathering information about children’s backgrounds through pre-event online sessions.

Some educators suggest children learn better through simple, emotionally gratifying answers, stories, songs, games, modern content, and interactive experiences, rather than complex-logical explanations or traditional lectures. They advocated adapting relaxed teaching styles with age-appropriate language, games, and multimedia, considering students’ backgrounds, interests, and prior knowledge (Fig. S3-9). A few educators emphasised direct, emotional experiences over lengthy lectures that move beyond traditional subject divisions.

*“Connecting concepts to real-world applications can enhance children’s understanding. I’ll ask them to crush a rock and observe the pores inside, and then submerge it in a bucket of water. Some water will disappear. Who drank it? If a small rock drinks this much water, how much this whole plateau can! - This way we can introduce porosity and permeability.”* - (E4; Male, 45 y.o.)

Educators also noted some changes to the sessions over the years. This mainly included children’s lessened interest in engagement, parent’s increased focus on the academic benefits, tightening of schedules (multi-day residential camps to single day events) and associated difficulties for educators. Educators preferred two-day residential camps over single-day events, contingent on the availability of logistical and parental support. They emphasised the unique first-time socio-emotional experiences of these camps, which help children, especially those who lack opportunities to form early positive associations with nature. They also believed these camps can foster appreciation, curiosity, and (sometimes strong) emotional connections to nature, and enable social interactions through cohabitation. In terms of content, a directed shift from experiences, observations and recreation to structured academic concepts was noted, according to some. Others said the content remained largely unchanged, but the focus gradually shifted away from quarrying issues.

> *“Earlier, children had ample free time to relax, explore, and interact with others. This schedule is very tight and too much for a day. Children do not get free time in between, and it is too academic. We only made it this way [perplexed smile].”* - (E10; Female, 40 y.o.)

While supporting the idea of initiating similar sessions much beyond Madayipara, educators observed challenges in incorporating diverse themes, such as history and culture, into these unfamiliar geographies. Many stressed the importance of documenting, evaluating, and updating sessions, particularly to avoid repetition among returning participants. Broader suggestions included sensitizing parents about the sessions and implementing policy changes to enable school teachers to conduct outdoor sessions. They observed a decline in school teachers’ enthusiasm for organizing similar events, attributing it to their limited interactive abilities, lack of geographical knowledge, and apprehension regarding vulnerability when engaging with children of diverse age groups. Recent policies on child safety also deterred school teachers from taking on responsibility for conducting the sessions due to limited logistical and manpower support.

Other general logistical challenges included financial constraints, reduced organizational strength and time, heavy tourism compromising the learning atmosphere, ensuring participants’ safety, unpredictable weather, and external resource person’s availability. Some suggested engaging public libraries and local sponsors as a solution to financial difficulties.

## DISCUSSION

Our quantitative results showed a short-term increase in overall knowledge, while there was considerable variation in knowledge gained across different themes. We suspect that these variations could be outcomes of the way the thematic sessions were implemented (e.g., themes that had more incorrect answers did not have much direct experiences associated with them). This finding and our pilot surveys suggests that quantitative measurements will largely depend on the specific topics that are discussed in these sessions, and importantly, formal knowledge gain can be varying among individuals, leading to 33-60% of participants showing no improvement. Additionally, conflicting ideas and confusion regarding conservation issues subsequently reflected in the open-ended responses may explain the wide variations in scores.

Qualitative analysis of educators’ and participants’ perspectives revealed major strengths and challenges of the sessions around the *Natureculture* elements, relational values, educators’ understanding of participants’ backgrounds and assumptions about children, pedagogical design, logistics, and awareness among parents and teachers. Bringing the qualitative and quantitative results in the light of existing literature, we identified some of the key insights emerging from a bio-culturally-rich, but neglected landscape. They are broadly concerning the content, logistics, session framework, educator’s approach and understanding of children’s background, and awareness for parents and teachers (Fig. 4). This broadly aligns with the framework of ‘Pedagogical Practices that Support Outdoor Learning Experiences’, centering on the learner, educator and the environment (Neville et al., 2023), but with more nuanced considerations for content and pedagogy.

**Figure 4.**
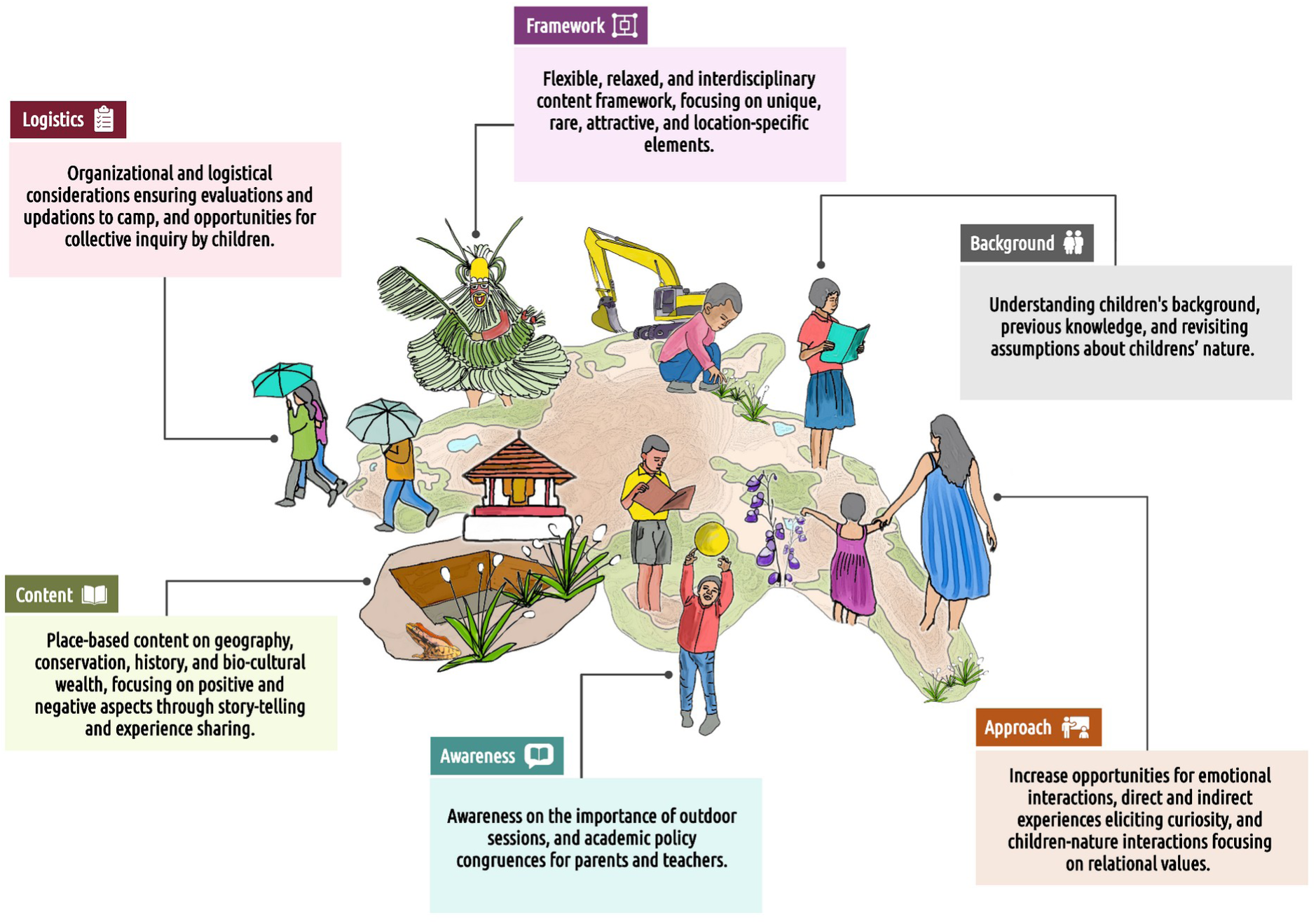
Major insights emerged from the perspectives of participants and educators of the conservation education sessions on rock outcrops of the Western Ghats. Representational figures indicating various elements used in the sessions on Madayipara, Kerala, India. Illustration by Jithin Vijayan.

### a. Children and educators on the session’s strengths

The sessions were able to provide knowledge on multiple aspects that are useful in curricular and extracurricular activities, and helped improve the multiple skills of children. It could also spread the messages of conservation. While clear aims about the camp worked well for some educators, others were more content with unstructured experiential learning. All educators’ aims are fulfilled in these sessions, and they agreed that these outdoor sessions can help children in forming positive associations with nature through unique and novel experiences, and catalyse more social interactions.

#### i. Unique experiences, *Natureculture* elements, and relational values

Participants noted several unique and engaging content and experiences in the sessions spanning from physical to human geography. Notably, participants focused on location-specific features, including rare and unique biodiversity, historical aspects, and aesthetics. These themes—cultural aspects, plateau uniqueness, ecosystem knowledge, and plant identification— were areas where children demonstrated strong understanding in the quantitative evaluation as well. More importantly, direct experience with nature (outdoor rain and cave experiences), interaction with peers, and exposure to new knowledge emerged as unique experience styles. Educators’ suggestion to incorporate game-based learning and outdoor explorations to engage with real-world problems, and use of narrative transportation to explain complex concepts aligns with previous research on how these can boost retention, persuasion, and attitude shifts (Ljungberg, 2024; Despland et al., 2025).

Participants’ and educator’s attachment to local heritage and frequently emerging *Natureculture* elements emphasize the importance of fostering a sense of relationship with heritage in conservation education (Richardson et al., 2025; Bouchard, 2025). It is also particularly interesting to note the emergence of rarely documented cultural ecosystem services such as cultural heritage (Gould et al., 2018). Participants also referenced loss of their ’native’ place, and peoples’ well-being – highlighting the importance of belonging (community) and identity, part of relational values (Gokhale et al., 2023; Chan et al., 2016). The increasing recognition that conservation perspectives are shaped by values, emotions, and experience— rather than solely by facts—is further supported by what children value. This aligns with the need for giving importance to contextual sociocultural values in conservation, to balance approaches that overemphasize the cognitive domain (Brias-Guinart et al., 2023; Toomey, 2023).

The reflected notions of cultural identity, well-being, and participants’ and educators’ desire for improved social cohesion underlines the importance of relational values in the conceptualization of nature conservation (Chan et al., 2016). These, along with the expressions of individual identity (place attachment) and social responsibility (care for children’s future) reiterates the potential power of relational values in strengthening the pedagogy. Without prompts through questionnaire items, multiple relational values emerged from children’s responses in interviews around responsibility, care, connectedness and stewardship, along with perceptions of identity and community (dos Santos & Gould, 2018). It was noted that ‘kinship’ did not reflect as a relational value in our study.

#### ii. Exposure on real-world conservation issues

The session conveyed rich information about environmental history, enriching children’s perspectives on protests, and rock outcrop conservation in a place-based way. Resulting diverse philosophical viewpoints on the intricate and multifaceted nature of conservation were consistent with earlier studies (Haydock & Srivastava, 2019; Spiteri, 2021). These perspectives included Deep Ecology (emphasizing habitat loss, species extinction, and cruelty to non-human animals), Ecological Modernization (positing compatibility between economic growth and environmental protection through technological and institutional reforms), impacts on human life, and the instrumental and relational values attributed to nature. This highlights the significance of fostering critical thinking, and encouraging autonomous and responsible decision-making on complex environmental challenges when educational programs are designed around real world issues (Varela-Losada et al., 2016; Kim et al., 2025).

### b. Children and educators on the session’s challenges

#### i. **Complexities of conservation communication**

The quantitative evaluation did not show any significant changes in attitudes towards environmental conservation, outdoor exploration, or the conservation of rocky outcrops. There was also a comparatively greater proportion of extreme increase and decrease in scores for identifying activities that could impact plateaus. These outcomes, coupled with the children’s simultaneous expression of hope and anxiety regarding the environment’s future and their perceived competence for action, warrants further investigation along with their meanings of nature conservation (Gokhale et al., 2023). Children pointing out the lack of ‘positive’ and ‘hopeful’ lessons, and doubting their competence in conservation action in our study, highlights the need for multiple strategies to help them cope with environmental changes (Chawla, 2020). Educators also acknowledged their ‘conservation vs. development’ dilemma influencing the sessions, which is documented earlier, and attributed to the intricate nature of conservation social issues (Ho & Seow, 2015; Israel, 2012). Taken together, this suggests that content and teaching styles could improve by adopting a more positive and constructive approach, and by considering both critical and abstract thinkers among the participants (Finnegan, 2023; Sutter et al., 2019). Educators can also employ appropriate conflict reflection tools, which can help them challenge their own perspectives and acknowledge the validity of diverse viewpoints and values (Hasslöf et al., 2014).

#### ii. **Need for harnessing the power of direct and unique experiences**

Participants’ were interested in observing organisms that are novel, unique, beautiful, rare, and curiosity-inducing. These characteristics, also identified in previous research (Fančovičová & Prokop, 2011; Prokop et al., 2025), will be valuable for future content development. Also, compared to plants, participants gained less knowledge about the animals, which could be due to relatively few direct experiences during the session, and perceived danger through peer discussions. This highlights the importance of addressing children’s geographies of fear in these sessions, giving equal importance to positive and negative nature experiences (Gokhale et al., 2023; Evans et al., 2023). Similarly, historical aspects and place knowledge were only slightly improved after the evaluated session, despite participants’ increased interest in related themes. This also needs further evaluation.

#### iii. Understanding the participants

Participants had diverse interests and sometimes contrasting viewpoints on how the sessions could be better. But in general, they desired content that was more relaxing, engaging, cross-disciplinary, and curiosity-inducing. Participants also wanted to explore the area on their own, with more guides, and opportunities to lead the sessions. Here, it is important to note that though pupil’s suggestions to teaching and learning activities tends to be very sensible, opportunities for these ‘student voices’ are rare in many cultures, and it might be challenging to educators to comprehend these (Skerritt et al., 2023). In our case, some educators already highlighted the challenges in understanding the backgrounds, strengths, and individual interests of all the participants, especially in multi-age-group sessions. Though challenging, effective conservation education programs necessitate a thorough understanding of participants’ backgrounds and interests; a critical insight highlighted in a prior global synthesis (Jacobson & McDuff, 1997). Thus, strategies such as pre-session activities to understand children (proposed by some educators in our study), and contemplative teaching practices might be needed to address these issues, strengthening the experiential learning cycle (Lee et al., 2020; Skerritt et al., 2023).

Educators also had contrasting views and assumptions on children’s learning capabilities, intentions for participation, ability for independent socialization, unguided explorations, and collective inquiry. These simplistic categories of children’s nature and their geographies in educators’ minds might need a critical evaluation, as children possess a multitude of identities that extend beyond these (Kaplan, 2024). This will help make design decisions for educational sessions, especially regarding the independence given to children, and educators’ facilitation roles (Thomas, 2010; Jamal et al., 2025). For example, it may be challenging to facilitate guided discovery learning and collective inquiry in children, granting children greater agency, as advocated by some educators and participants. But this can be done by adapting existing designs from other contexts (Janssen et al., 2014) to local settings in a collaborative effort involving parents, organizers, and teacher trainees, with due consideration for safety.

Educators and participants also emphasized connecting new information to prior knowledge, highlighting the importance of scaffolding that bridges classroom learning with real-world application to ensure memorable, comprehensive, and long-term learning (James & Williams, 2017). Some educators’ thoughts also aligned with previous studies, which highlighted the importance of positive psychology, contemplative practice, storytelling, emotions, individual experiences, interactions, and slow, cross-disciplinary pedagogies in outdoor learning curriculums to build relational experiences with place (Gray & Pigott, 2018; Hunter & Campbell, 2025). Also, educators highlighting the need for ‘emotional’ experiences over ‘logical’ content in our study, emphasizes the integration of socio-emotional learning frameworks with ideas of social-emotional functioning, cognition, motivation, and learning (Immordino-Yang et al., 2019; Prakash, 2023).

#### iv. The implementation challenges

Our findings underscore varied implementation challenges, consistent with previous studies (Yemini et al., 2023; Mann et al., 2022b). These include limitations in funding and time, logistics, lack of evaluations, a crowded curriculum, insufficient teacher training in student-centred and outdoor pedagogies, ensuring participants’ safety, and the demands of standards-based learning. Raising awareness among parents and teachers about the contributions of these sessions is also challenging (Kuo et al., 2019). Policy changes are also necessary to ensure teachers implement such sessions in schools, rather than merely suggesting them in the curriculum. Given the significant role of NGOs in implementing EE across the globe, their long-term expertise, and insights, as documented in this study and existing frameworks will be valuable in addressing these challenges (Nature Classrooms, 2021; Alam, 2023).

### c. Limitations of the study

While efforts were made to address the issues of simplistic design and inadequate controls (Miller et al., 2021) in our study, the current design was inevitable considering the logistical feasibility. Despite our best efforts, children might have mimicked their peers when responding to survey questions, and they may have been less familiar or comfortable with questions about their attitudes and perceptions. To address this, future long-term research can employ robust quantitative instruments to understand changes in attitudes and behavioural outcomes, which are not yet locally-validated for similar contexts (Powell et al., 2019), and are relatively long, considering children’s fatigue, and time availability.

## CONCLUSION

The long-term experiences of grassroots initiatives explored in this study provide valuable insights for understanding the nuances in place-based education (Hill et al., 2025). Further long-term socio-ecological assessments of these initiatives can offer insights for neglected conservation education contexts in the global South (Ardoin et al., 2020; Nielsen et al., 2021). Our study highlights the importance of emotions, positive and negative experiences, positive perspectives, relational values, and contemplative pedagogy when designing conservation education sessions, while critically examining educators’ assumptions about children’s nature and geographies. In addition to the insights from educational and psychological aspects, our insights also underline the advantages to content design when they are contextualized in multifaceted socio-ecological systems. Complementing the existing designing and assessment frameworks, they provide nuanced perspectives on content and pedagogical considerations useful in often overlooked socio-ecological contexts.

## ACKNOWLEDGEMENTS

Our sincere thanks to all parents, teachers, children, and residents of our study areas for their wholehearted cooperation and support during the fieldwork. Our heartfelt thanks to Suhel Quader, Ovee Thorat, and Aparna Watve for their extensive help during the study conceptualization, designing, analysis, and for comments on earlier versions of this manuscript. We thank Ricardo Correia for his comments that improved this manuscript. We thank members of Kerala Sasthra Sahithya Parishad (Madayi region), Society for Environmental Education in Kerala and Malabar Natural History Society; Anaswar K, Anjitha Vijayan, Bhargavan PK, Ganeshan, Haridasan Naduvalath, Prasad PV, Rajeswari Bhai BT, Ratheesh C, Sheeja Vijayan, Suri Venkatachalam, Vena Kapoor, Afna P, Yathumon MA, and Yuvan Aves for their extensive support and encouragement during the study. We thank On the Edge Conservation (UK) for funding this work, and Nature Conservation Foundation (India) team for the logistical support. VJ thanks the organizers and participants of the workshop ‘Perspectives on Learning’ by Digantar Shiksha Evam Khelkud Samiti and Wipro Foundation for insightful learnings.

## FUNDING

On the Edge Conservation, UK.

## AUTHOR CONTRIBUTIONS

Vijayan Jithin conceived the ideas and designed the methodology with inputs from Rohit Naniwadekar. Vijayan Jithin collected, analysed, and led the writing of the manuscript. Both authors contributed critically to the drafts and gave final approval for publication. Our research was discussed with local stakeholders at multiple stages, to seek feedback on the questions to be tackled, and the approach to be considered. Literature published by scientists and local researchers from the region are cited where relevant; and we have considered relevant work published in the local language, Malayalam.

## COMPETING INTERESTS

Authors declare that they have no known competing interests.

## DATA AND MATERIAL AVAILABILITY

We are unable to make data on participants’ responses to the interviews publicly available, as the transcripts contain sensitive information that cannot be fully anonymised. All codes used in the analysis will be made available in Zenodo after acceptance, along with anonymized data from the written survey responses.

## Notes

### Competing Interest Statement

The authors have declared no competing interest.

