## Supplementary material for "Listening to children’s and educators’ experiences in long-term place-based conservation education sessions on southern Indian rocky outcrops": Fig. S

**FIGURE S1**
Violin plot showing the changes in participants' overall plateau knowledge score before and after the field trip session.

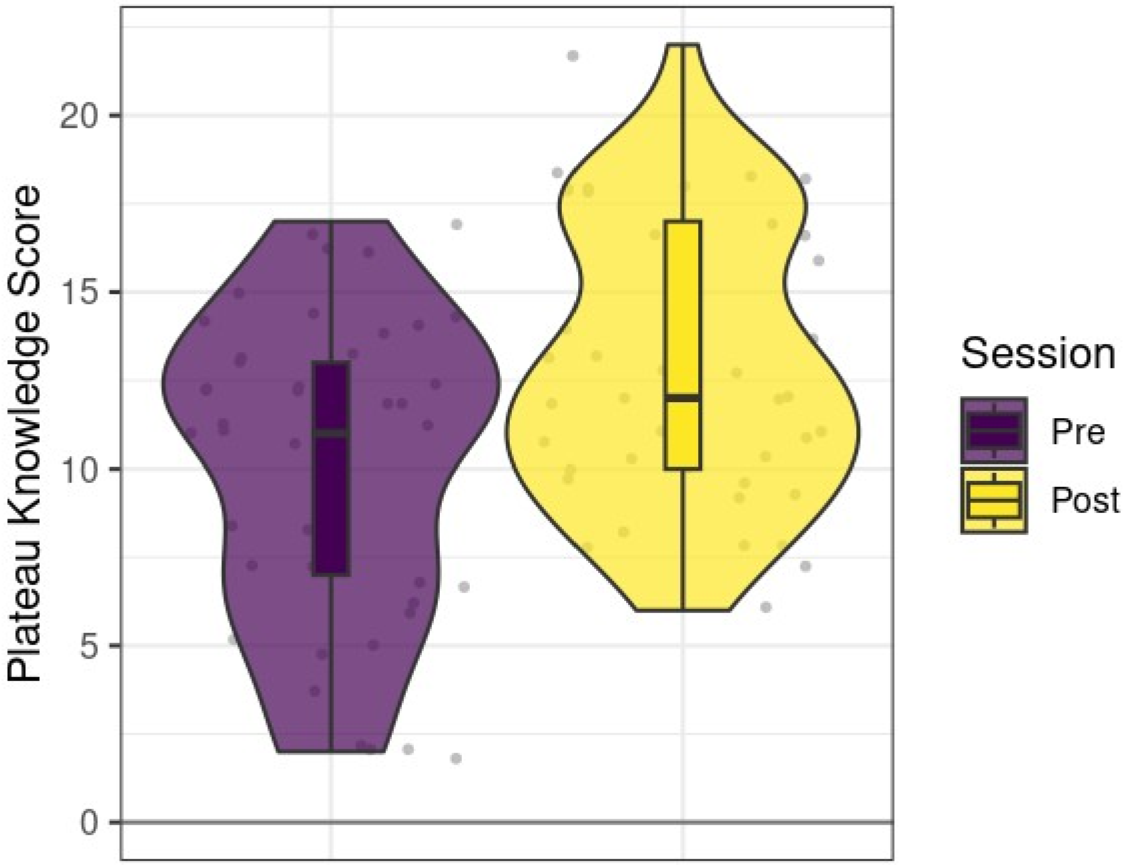

**FIGURE S2**

A. Participants' response to the question (a) '*why do you think these activities will alter the current state of plateaus?*', and (b) '*what are the first five thoughts coming to your mind when seeing this news?'* [article about people protesting against quarries in Madayipara].

(a)

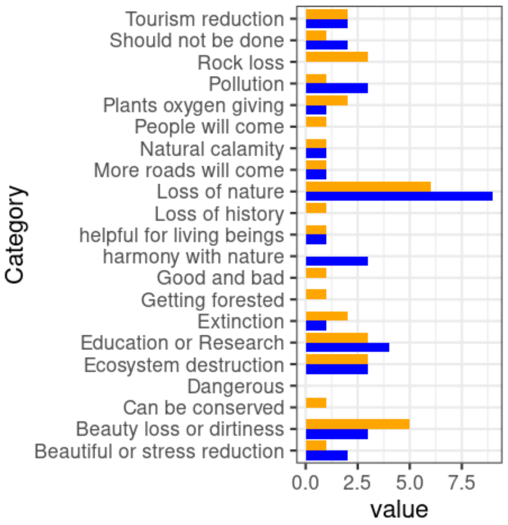


(b)

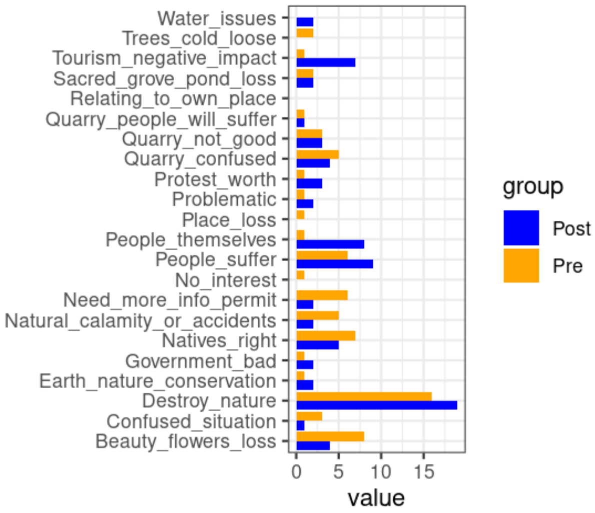


B. Participants' reasonings for the interest or disinterest in seeing Scorpion and *Drosera*.

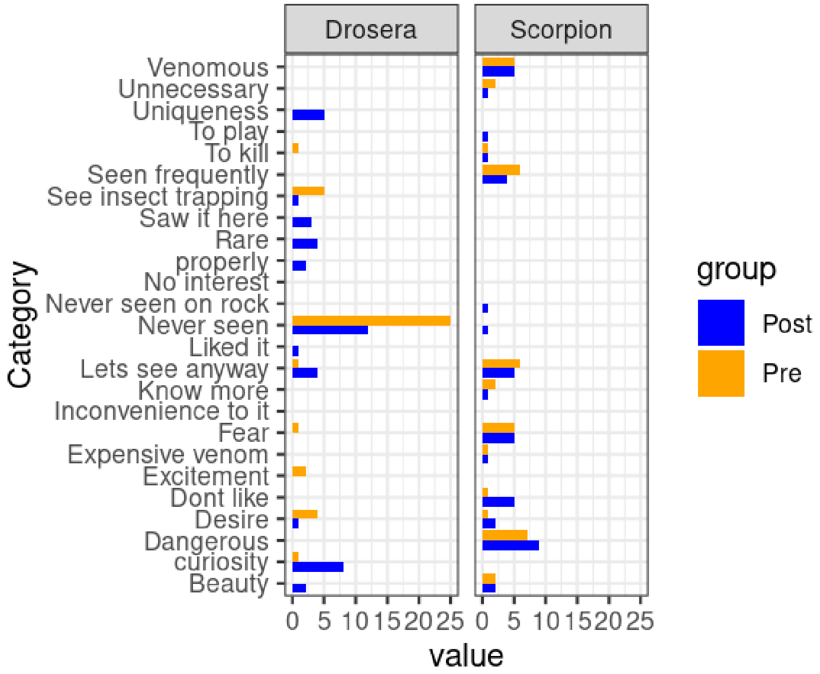

**FIGURE S3**

Hierarchical nature of themes and sub-themes emerged in the qualitative analysis, along with some relevant quotes from the interviews.

1. **Participants' knowledge**

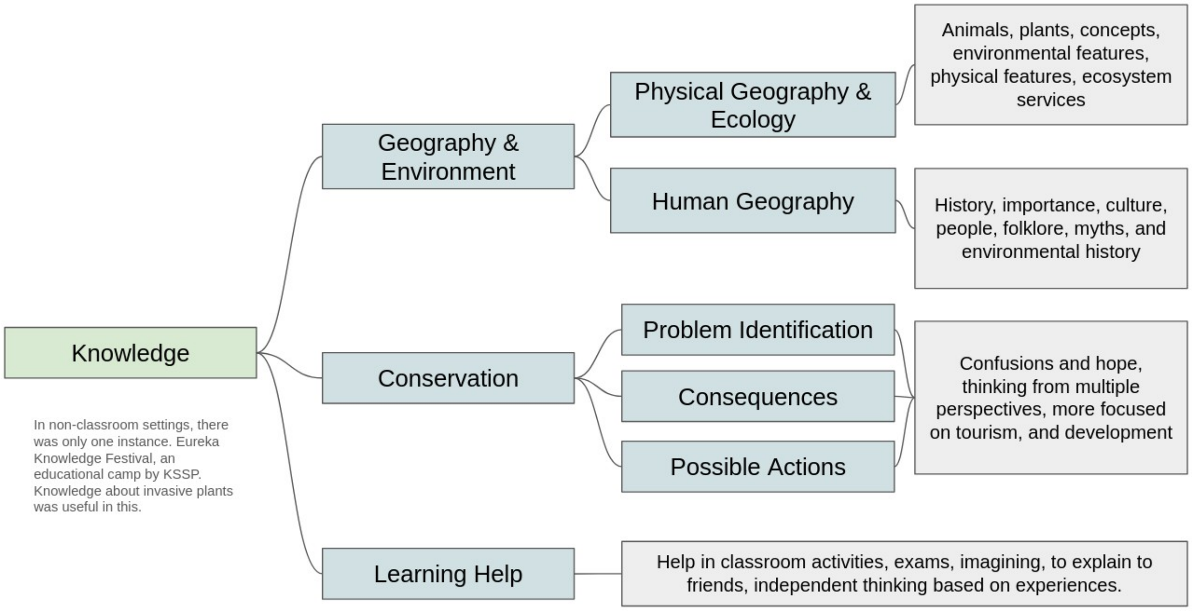


2. **Participants' experience**

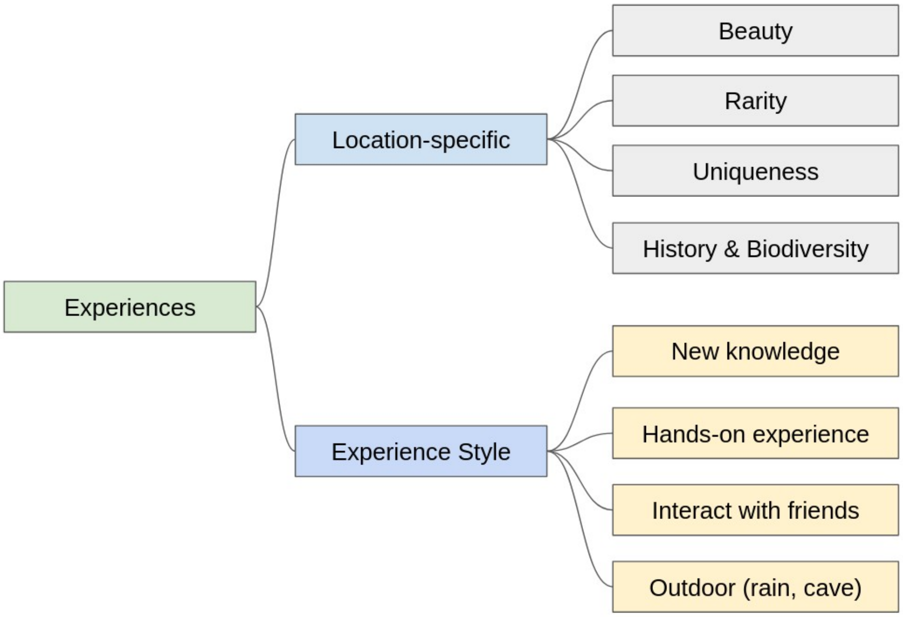


*"It is a place where we can go and spend time.. Observe nature..How much ever stress you have, you can go sit there, you'll feel relaxed"* -  - (P8; Male, 16 y.o.)

#### 3. Participants' suggestions

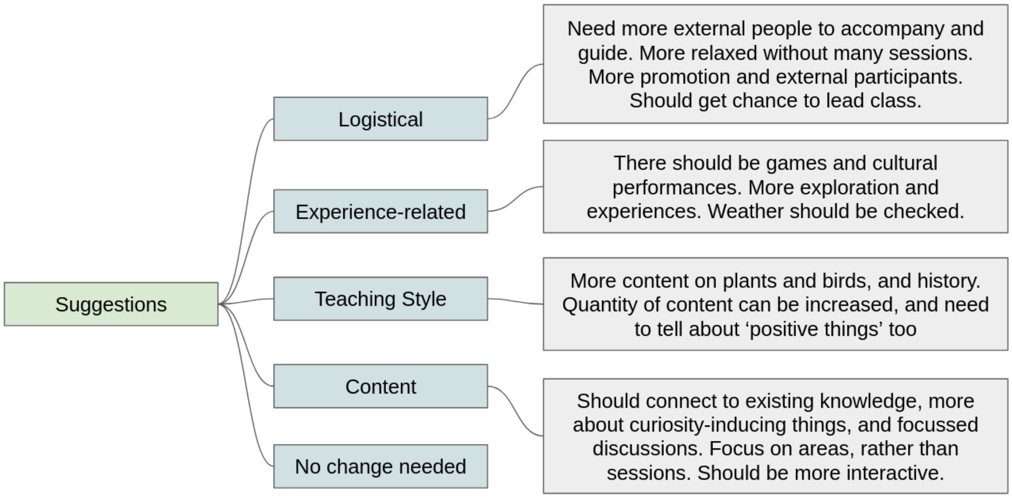


*"So many things were told about the exploitation of nature, and all, right? There need to be more 'good things' about Madayipara and nature. Also, we need to increase the number of children and elders participating together. That day everyone should keep their mobile phones and other things aside"* - (P3; Male, 12 y.o.)

*"Instead of classifying it [the session] into plants, animals etc.. We can focus on places where we are, and then talk about multiple things there."* - (P10; Female, 11 y.o.)

#### 4. Educators' aims

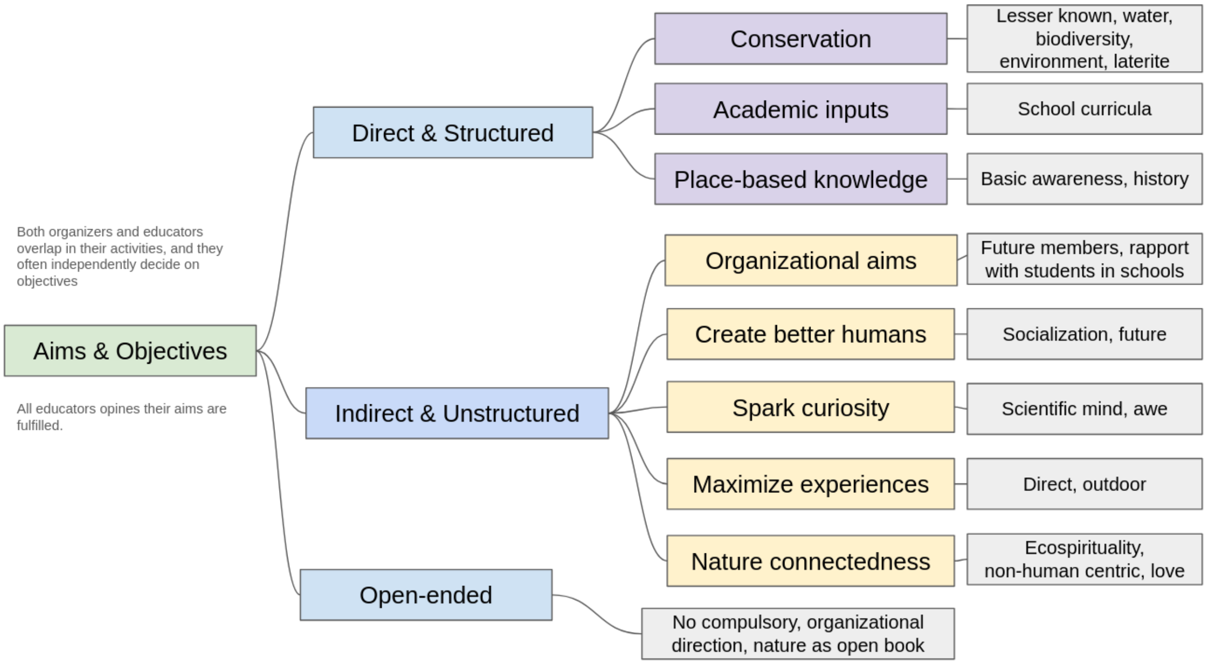


Note: The indirect aims also include attracting new members to host organizations, and build student rapport in schools.

### 5. Educators' motivation

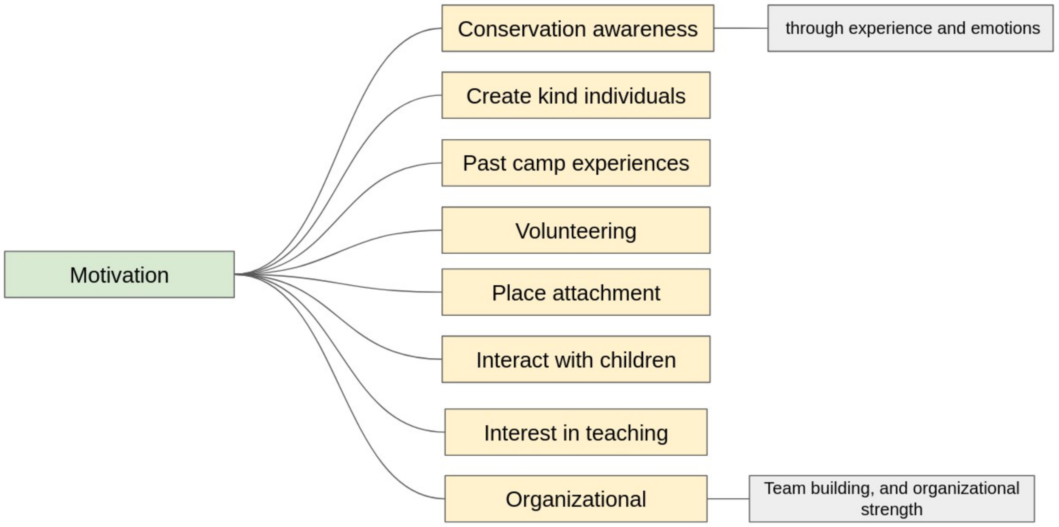


### 6. Educators' preparations

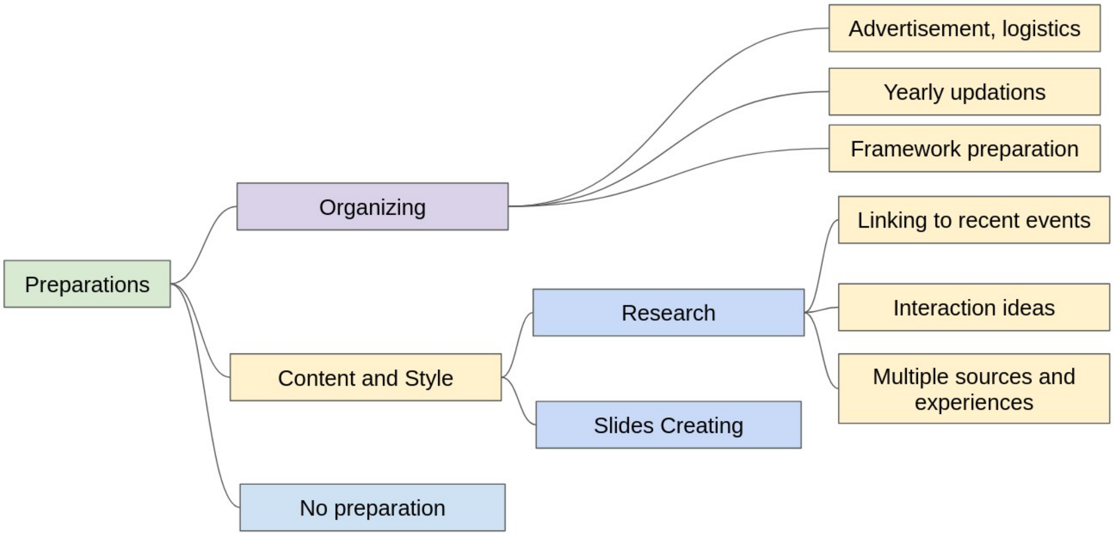


Preparations mainly included making logistical arrangements, promotions, making yearly updates, and finalizing the overall framework. Some educators prioritized developing or collating ideas for interacting with children, finding contemporary content from recent children's literature, and devising relevant games.

> *"...On the [first] day, we can present a 1-1.30 hrs long slideshow of different things that can be seen the next day, and stories about them. Where to find and how to identify them. Next day, children will explore the area without field guides, and ask us "This is that one, no?"; going with pre-defined aims of first seeing, and then to check if what was told yesterday is correct or not [smiles]. It is fun for them. They'll discuss that with each other. For children, all adults in the camp are guides for them. Whoever it is; it could be a parent as well. Which means we need a lot of such 'field guides' to cover that large rock expanse."* - (E2; Male, 49 y.o.)

7. **Educators' view on the changes over the years**

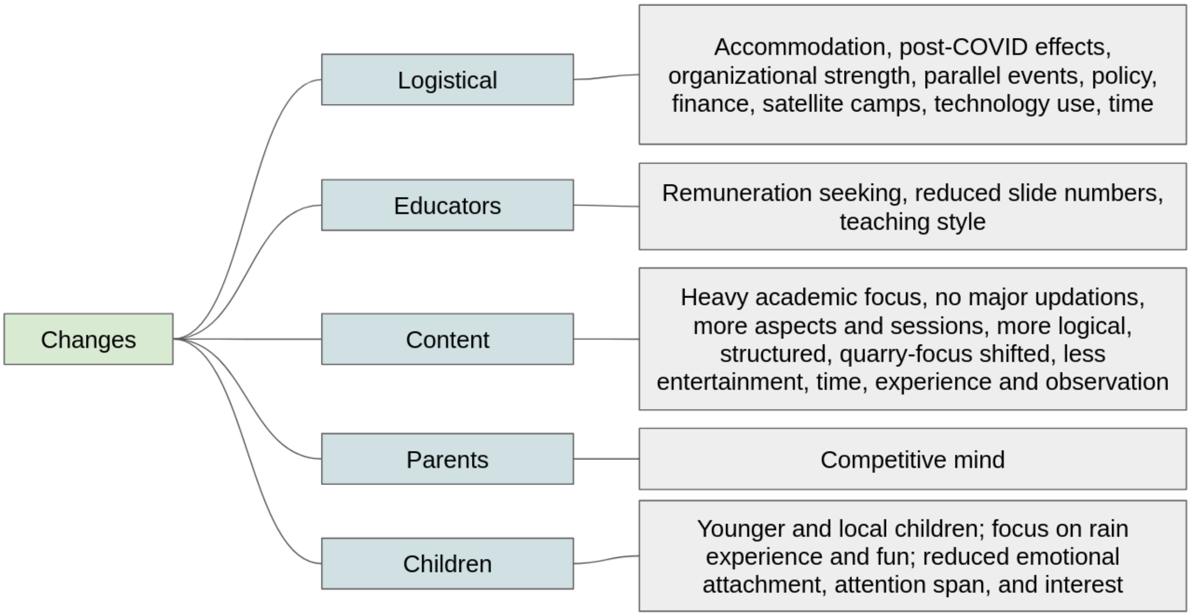


Educators noted changes to the sessions from multiple perspectives. Over the years, some sessions have focused on encouraging participation of younger children from nearby areas. According to the educators, participants over the last few years were less interested in the content, with limited attention span and emotional attachments, and more interested in the rain experience and recreation. Educators also reported that, increasingly, the motivation of the parents to send children were the potential benefits of the camp to the academic prowess of their children. Educators were also changing teaching styles to match the tight schedules. While earlier the educators conducted the sessions voluntarily, recently some educators have started seeking remunerations. Educators also noted the emergence of parallel satellite events on plateaus other than Madayipara and increased use of multimedia over time. Some multi-day residential camps are now conducted as single-day events, thereby resulting in tighter schedules. This is mainly due to reduced organizational strength, financial constraints, and COVID-19 pandemic.

### 8. Educators' challenges

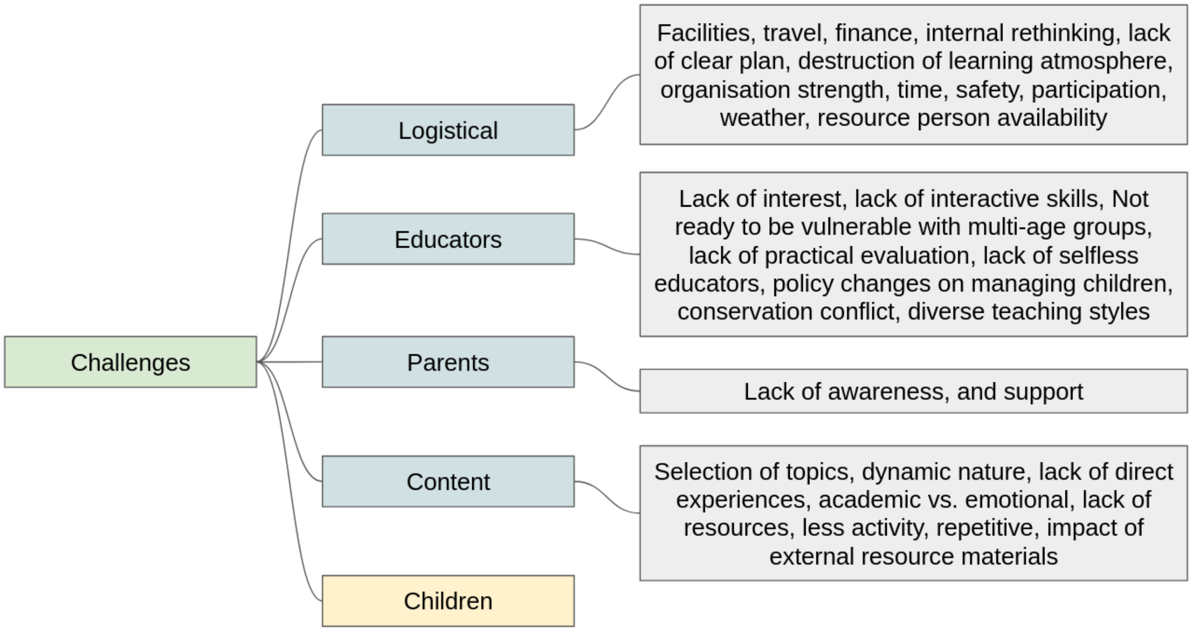


*Much more nuanced, and detailed; see below:*

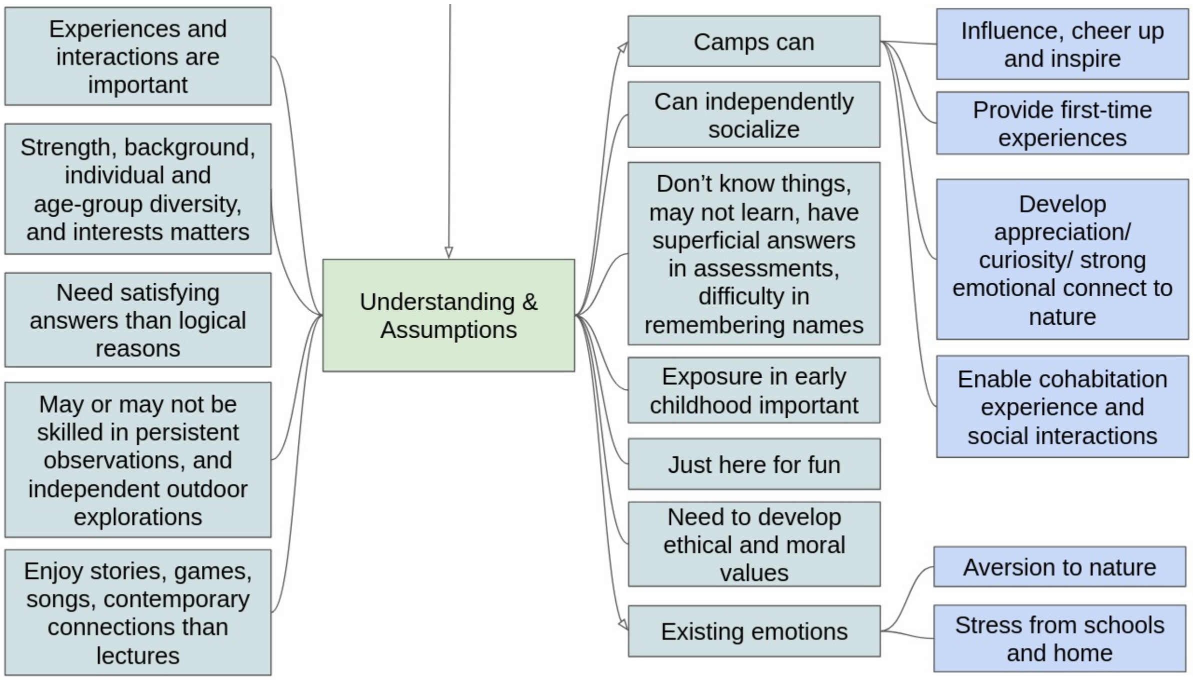


*"In one camp, there was a child who only collected pebbles. He was not interested in plants or dragonflies or anything. He asked me to put all those in my bag. He was carefully collecting and classifying the pebbles from there. He was so keen on that. Some inspiration from within him. Also there were children who only looked for birds, some focusing on the interaction between birds and dogs; they are always behind that. They'll get involved in their own interests, and they will find their own new worlds from the connections that they made."* - (E2; Male, 49 y.o.)

### 9. Educators' suggestions

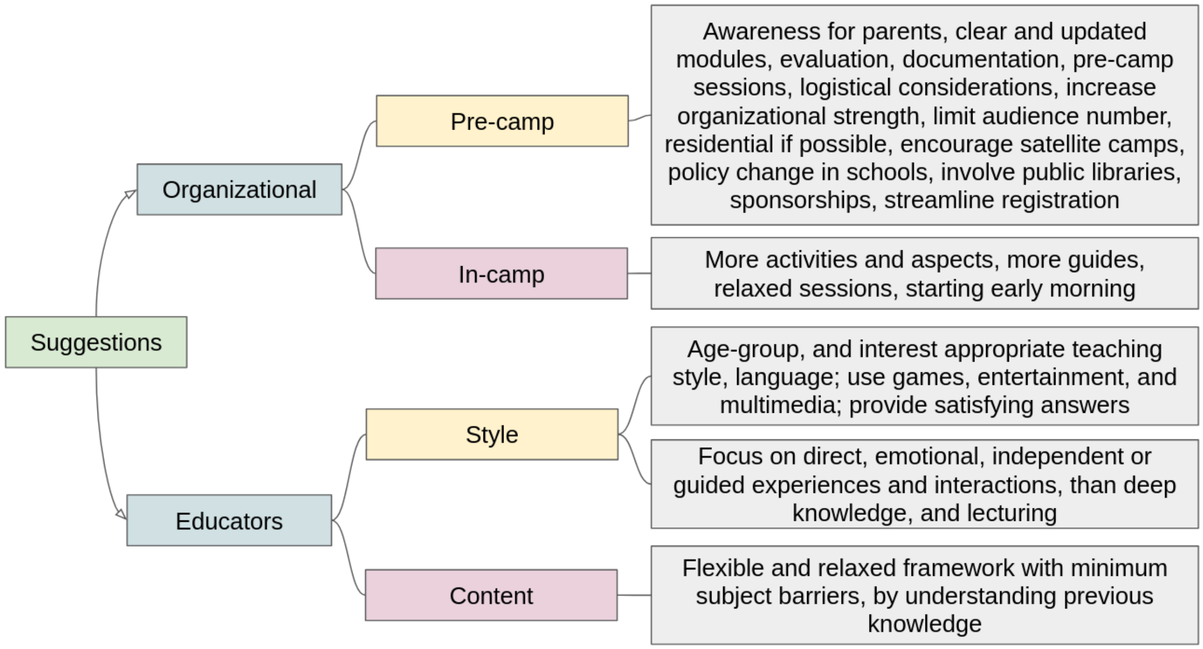


*"We need a romantic approach here. Just dry statistics or science won't suffice. We can present such stories and poems [referring to the ones described earlier in the interview]. Humans should feel pain when they see a life is no more, no?. The American writer Norman Cousins says that 'The first aim of education should not be to prepare young people for careers, but to enable them to develop respect for life"* - (E1; Male, 67 y.o.)

*"It will be better if it becomes [again] residential. The evening time feeling, the 'vibe' that today's children talk about is there during that time. It is okay even if it is one day, but it should conclude by 8 pm..But organising it will be an issue..or we should utilise the early morning time. Madayipara is particularly beautiful in the evening. Arranging homestay experience will be also great. I have done similar two day camps in school"* - (E13; Male, 49 y.o.)

**TABLE S1**

Table showing the outputs from Cumulative Link Mixed Models for attitude-related statements. Models fitted with Laplace approximation, and ‘probit’ links.

| **Statement** | **Mean** | **SE** | ***z*** | ***p*** | **Random effect (ID) variance** | **SD (ID)** | **N** |
|---|---|---|---|---|---|---|---|
| I think this habitat, and organisms living in this habitat do not need any special protection | 0.1765 | 0.2474 | 0.714 | 0.475 | 0.6632 | 0.8144 | 41 |
| We can solve environmental problems | 0.2949 | 0.2547 | 1.158 | 0.247 | 0.9907 | 0.9954 | 41 |
| Rock outcrops are officially classified as ‘wastelands’. I think this is appropriate | 0.2501 | 0.2440 | 1.025 | 0.305 | 0.4765 | 0.6903 | 40 |
| I like to watch plants and animals and observe my surroundings | -0.2530 | 0.2807 | -0.902 | 0.367 | 0.5414 | 0.7358 | 41 |

**APPENDIX S1**

This camp included four sub-sessions on plants, animals, conservation, and history of the area by different educators and lasted between 9:00 AM and 6:00 PM. For this, following a brief introduction about the study to both parents and children, the pre-camp survey was administered to all participants guided by the researcher, by reading aloud each item and giving students sufficient time to answer (Larson et al., 2011). A post-camp survey that contained the same questions, and a few additional questions on the camp experience was conducted at the end of the camp. For attitude related questions, a five-point Likert Scale response system (strongly disagree, disagree, no opinion/ not sure, agree, strongly agree) was used with facial emojis along with text in *Malayalam* (Massey, 2022). Questions were designed after appropriate assessments for directionality. Participants of the pilot-test were aged between 9-15, and from nearby Panchayats. After analysing the results of this pilot-test considering the notes made by the researcher during the camp, and after seeking feedback on language-use, and comprehensibility from a few participants, we refined our questionnaire with more open-ended questions (Jithin & Naniwadekar, 2025) to capture the direction of and variation in the attitudes (Salazar et al., 2021).

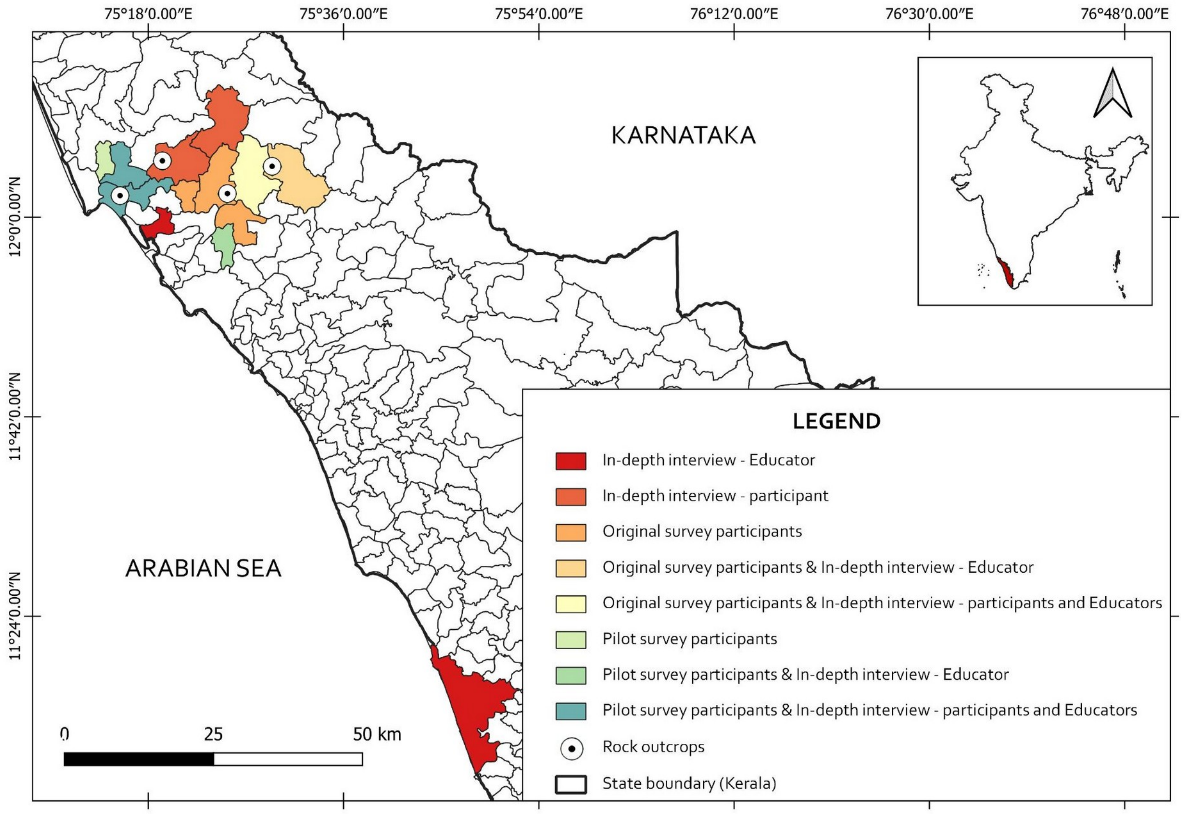


Map showing the locations of focal rocky outcrops covered in this study, and Gram Panchayats (basic governing institutions in Indian villages, representing groups of villages) from where participants and educators for the surveys and in-depth interviews are coming from (Kerala, India). The number of samples from each Panchayats and names of the outcrops are removed to ensure anonymity of individuals and the educational institutions involved.


**Questionnaires:**

Pilot Test Questionnaire and Revised Questionnaire in *English* and *Malayalam*:
https://doi.org/10.5281/zenodo.15868707

**APPENDIX S2**

Demographic Information about the educator interviewees. Detailed information and other backgrounds are not included to ensure anonymity. Note that serial numbers are assigned randomly, and the participant number given along with the quotes are not matched with these.

| Sl. No. | Gender | Age | Experience and background |
|---|---|---|---|
| 1 | Female | 40 | Upper Primary School Teacher |
| 2 | Female | 56 | NGO Member, Retired Primary School Teacher |
| 3 | Male | 44 | Higher Secondary School Teacher, NGO Member |
| 4 | Male | 64 | NGO Member, Organizer of long-term educational sessions |
| 5 | Male | 49 | Higher Secondary School Teacher, NGO Member |
| 6 | Male | 49 | Higher Secondary School Teacher, Teachers' Trainer, NGO Member |
| 7 | Male | 52 | Upper Primary School Teacher, NGO Member |
| 8 | Male | 55 | Upper Primary School Teacher, Curriculum Committee Member, NGO Member |
| 9 | Male | 67 | Naturalist, Educator, NGO Member |
| 10 | Male | 41 | High School Teacher |
| 11 | Male | 45 | University Professor, NGO Member, Educator |
| 12 | Male | 56 | Scientist, NGO Member, Educator |
| 13 | Male | 65 | University Professor, Educator |
| 14 | Male | 54 | University Professor, Educator, Ecologist |
| 15 | Female | 63 | Retired Primary School Teacher, NGO Member |
